# Is the addition of higher-order interactions in ecological models increasing the understanding of ecological dynamics?

**DOI:** 10.1101/595140

**Authors:** Mohammad AlAdwani, Serguei Saavedra

**Affiliations:** Department of Civil and Environmental Engineering, MIT 77 Massachusetts Av., 02139 Cambridge, MA, USA

**Keywords:** Lotka-Volterra models, Free-equilibrium points, Bernshtein’s theorem, Polynomial dynamical systems, Explanatory power

## Abstract

Recent work has shown that higher-order interactions can increase the stability, promote the diversity, and better explain the dynamics of ecological communities. Yet, it remains unclear whether the perceived benefits of adding higher-order terms into population dynamics models come from fundamental principles or a simple mathematical advantage given by the nature of multivariate polynomials. Here, we develop a general method to quantify the mathematical advantage of adding higher-order interactions in ecological models based on the number of *free*-equilibrium points that can be engineered in a system (i.e., equilibria that can be feasible or unfeasible by tunning model parameters). We apply this method to calculate the number of free-equilibrium points in Lotka-Volterra dynamics. While it is known that Lotka-Volterra models without higher-order interactions only have one free-equilibrium point regardless of the number of parameters, we find that by adding higher-order terms this number increases exponentially with the dimension of the system. Our results suggest that while adding higher-order interactions in ecological models may be good for prediction purposes, they cannot provide additional explanatory power of ecological dynamics if model parameters are not ecologically restricted.

## Introduction

Lotka-Volterra (LV) models (Lotka, 1920; Volterra & Brelot, 1931) have provided fundamental insights about ecological systems for almost a century (Case, 2000). Yet, it is known that LV models are parsimonious approximations (MacArthur, 1970; Strogatz, 2015) and do not capture all the complexities arising from the dynamics of ecological systems under investigation (Hairston et al., 1969; Momeni et al., 2017). Many times, the prediction errors of LV models have been attributed to the existence of higher-order interactions (HOI) (Abrams, 1983). More broadly, HOIs can be seen as the “unseen” influences of one species on the others. That is, the effect of species A on the per capita growth rate of species B might itself depend on the abundance of a third species C due to either compensatory effects or supra-additivity (Case, 2000). Therefore, HOIs have been typically translated as addition or modifications of higher-order terms in existing population dynamics models (Case & Bender, 1981).

It has been shown that HOIs can stabilize dynamics in competition systems (Grilli et al., 2017), promote diversity in ecological communities (Bairey et al., 2016), capture unexplained complexity of LV models (Friedman et al., 2017; Mayfield & Stouffer, 2017), and dominate the functional landscape of microbial communities (Sanchez-Gorostiaga et al., 2018) (but it has been shown that HOIs play a non-significant role in predicting protozoan populations, Vandermeer 1969). However, it remains unclear whether these apparent explanatory benefits of adding higher-order terms into population dynamics models come from fundamental principles or from simply exploiting a mathematical advantage given by the nature of multivariate polynomials. Yet, answering this question is important in order to increase our understanding about how to best investigate the role of higher-order interactions in shaping ecological dynamics.

In general, it is expected that adding HOIs into population dynamics models, the number of parameters increases, facilitating the capacity of the model to fit experimental data or empirical observations. While studies have been statistically penalizing for this increase (Aho et al., 2014; Mayfield & Stouffer, 2017), it is still unclear whether the number of parameters is the main factor controlling the degrees of freedom of ecological models. For example, the existence of feasible equilibrium solutions is a crucial condition in the context of species coexistence in ecological dynamics of the form *dN*_*i*_/*dt* = *N*_*i*_*f*(***N***) (i.e., a necessary condition for persistence, permanence, and the existence of bounded orbits in the feasible domain, see Hofbauer & Sigmund 1998; Song & Saavedra 2018; Stadler & Happel 1993). Yet, two dynamical models with the same number of parameters can have different numbers of free-equilibrium points (i.e., solutions that can be either feasible or unfeasible by tunning model parameters). Note that it is possible to predict either species coexistence or non-coexistence by engineering free-equilibrium points into feasible or unfeasible solutions, respectively. In the classic LV model, there is only one free-equilibrium point regardless of the dimension of the system (Takeuchi, 1996). However, it is unclear the extent to which HOIs can increase the number of free-equilibrium points and multiply the ways of reaching any ecological dynamics, which standard statistical methods cannot penalize for (Akaike, 1974; Schwarz, 1978). In other words, is the explanatory power of HOIs a mathematical construct that comes from feeding more free-equilibrium points into ecological models?

To answer the question above, first we investigate the effect of adding free-equilibrium points on the capacity to engineer the behavior of ecological systems. Specifically, we illustrate this effect through a simple example, which shows that by adding a single free-equilibrium point to a 2-species system, one can more easily achieve both feasible and unfeasible solutions compared to the LV model. Then, we introduce a general method, based on Bernshtein’s theorem (Bernshtein, 1975), to count the number of free-equilibrium points of polynomial dynamical systems. Then, we apply this method to show that the number of free-equilibrium points in LV models with HOIs increases exponentially with the dimension of the system, inheriting the engineering advantage of polynomial dynamical systems. Finally, we discuss the implications of our findings in the context of fitting HOIs to empirical data and the use of higher-order terms in general.

### Understanding the effects of HOIs

To understand the effects of adding HOIs on the dynamics of ecological systems, we considered the simplest possible LV model with HOI terms. This model involves the classic 2-species LV system with an additional non-additive interaction term per capita *N*_1_*N*_2_:

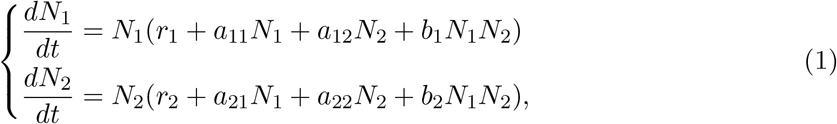

where *N*_*i*_ and *r*_*i*_ correspond to the abundance (biomass) and maximum per capita growth rate of species *i*, respectively. Additionally, *a*_*ii*_ corresponds to self-regulation terms, *a*_*ij*_ corresponds to interspecific terms, and *b*_*i*_ corresponds to HOI terms.

In general, it is known that the equilibrium points of a system like Eqn. (1) are given by the intersection points of the systems’ isoclines, which are obtained by setting the time-derivatives to zero (Strogatz, 2015). We can classify these equilibrium points into either *rigid* or *free*. We defined rigid-equilibrium points to be the ones restricted to particular subsurfaces of the space regardless of the values that the model parameters can take. Hence, there is less flexibility in terms of controlling their locations in space. For example, in the classic 2-species LV model (without HOI terms, i.e., *b*_1_ = *b*_2_ = 0 in Eqn. 1), no matter how we change the model parameters, one equilibrium point will be always at the origin, while two equilibrium points will lie always along each of the axes. Thus, in the LV model, rigid-equilibrium points can be defined as the ones which contain at least one zero coordinate (i.e., boundary-equilibrium points). On the other hand, we defined free-equilibrium points as the ones whose locations are not restricted in space and are completely dependent on model parameters. As mentioned before, it has already been proved that LV models without HOI terms have one single free-equilibrium point (Takeuchi, 1996).

While previous studies (Berg, 2000; Biroli et al., 2018; Galla, 2007) have focused on estimating numerically the total number of equilibrium points (i.e., without separating rigid- from free-equilibrium points), only free-equilibrium points dictate the dynamics of a feasible system (Song et al., 2018). As we mentioned before, the existence of a feasible solution is a necessary condition for persistence and permanence in dynamical models of the form *dN*_*i*_/*dt* = *N*_*i*_*f*(***N***) (Hofbauer & Sigmund, 1998; Stadler & Happel, 1993). Similarly, it has been proved that this type of models cannot have bounded orbits in the feasible domain without a feasible free-equilibrium point (Hofbauer & Sigmund, 1998). In fact, due to the non-revival property of such models (Takeuchi, 1996), the rigid-equilibrium points are the free-equilibrium points of the same model after substituting the corresponding zero abundances (of the species that die out) and deleting the equations which involve their time derivatives (as they will be zero as well). Therefore, for the purposes of this work, we will focus on the free-equilibrium points of LV models. Yet, for the interested reader, we have derived an analytic formula that depends solely on the number of free-equilibrium points to count the total number of equilibrium points in a LV model with or without HOI terms (see Supplementary Information).

To obtain the free-equilibrium point(s) of a system and its (their) effect on the dynamics of a system, we need to obtain the *free*-isocline equations, which are the classic isocline equations but considering that no state variable can take a value of zero. For example, the free-isocline equations for system (1) read:

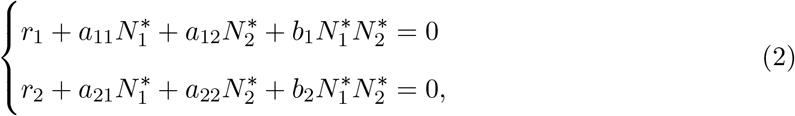

where 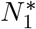 and 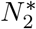 are the steady-state abundances. Then, to provide the number of free-equilibrium points for this system, we can rewrite Eqns. (2) such that each equation is expressed in terms of a single state variable:

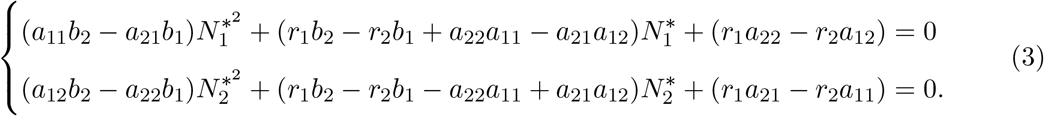

The uncoupled Eqns. (3) can be solved independently via the quadratic formula. We denote the solutions of the first and second equations by 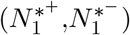 and by 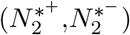, respectively. Note that 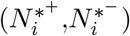 are the solutions to the quadratic equation where the sign of the square-root of the determinant is positive and negative, respectively. It is easy to check that 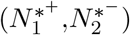 and 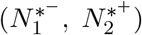 are the solutions to Eqns. (2). These solutions reveal that system (1) has 2 free-equilibrium points.

Note that if *b*_1_ = *b*_2_ = 0, the two separate solutions 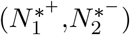 and 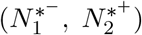 collapse into a single one (i.e., 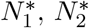). Under this condition (i.e., the classic 2-species LV model without HOI terms), species will coexist when the only free-equilibrium point is both feasible (i.e., positive) and stable (Saavedra et al., 2017). However, if *b*_1_ and *b*_2_ are not zeros, species coexistence will be attained when at least one free-equilibrium point (out of the 2) is feasible and keeps the trajectory of the initial condition close to it at all times. That is, the existence of the extra free-equilibrium point makes it easier to bring at least one free-equilibrium point to the feasible domain via parameter tuning (if no parameter restrictions are imposed a priori).

Hence, because of the flexibility that is introduced via additional free-equilibrium points, one may be tempted to conclude that HOI terms do promote species coexistence (or feasible solutions). However, it is easy to show that the conditions to achieve non-coexistence (or unfeasible solutions) also increase by adding HOIs. For instance, apart from manipulating the species abundances in the real domain, we can impose that any one of the following quantities 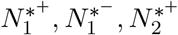, or 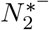 has an imaginary component, making both solutions 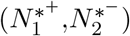 and 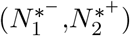 outside the feasible domain (due to the quadratic formulation of the uncoupled Eqns. (3), imposing 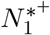 to have an imaginary component implies that its complex conjugate in the other solution tuple 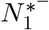 has an imaginary component as well). Note that engineering imaginary steady-state abundances from real model parameters is impossible to attain in the classic LV model without HOIs (Takeuchi, 1996). Therefore, HOIs increase the flexibility of the system to achieve either coexistence or non-coexistence (given that the imaginary domain for abundances has become accessible).

Finally, it is important to notice that the overall flexibility gained by adding extra HOI terms can further increase depending on the model. For example, by adding quadratic terms per capita (i.e., 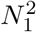 and 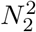) to Eqn. (1), it can be shown that the system will have 4 free-equilibrium points and 4 sets of separate solution tuples, from which we need to calibrate at least one of them to achieve coexistence (or manipulate the abundances of a few species in all tuples to achieve non-coexistence). That is, HOIs can increase the number of free-equilibrium points, and in turn, increase the flexibility to arrive to any possible conclusion. In the context of fitting (Mayfield & Stouffer, 2017), while the free-equilibrium points can be feasible or unfeasible, without parameter constraints these free-equilibrium points can always be engineered to find feasible solutions. For example, in the analysis of time-dependent quantities (such as species abundances), the most typical approach is to use the initial time points as the initial conditions to fit the dynamics of a system. Unfortunately, this initialization already biases and engineers the solution (Abu-Mostafa et al., 2012), which is relatively easier to achieve with higher-order terms. Therefore, the number of free-equilibrium points is a measure of how easy it is to fit data to a dynamical model (which is not necessarily linked to the number of parameters or species), but not necessarily about the ecological mechanisms of a system. This is a key problem we turn our attention in the next section.

## Methods

### Quantifying the effects of HOIs

To introduce a general methodology to quantify the number of free-equilibrium points in ecological models, we used a generic system of polynomial dynamical equations for *n* species of the form:

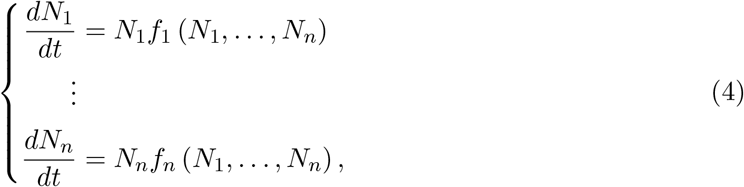

where *f*_1_, *f*_2_,…, *f*_*n*_ are multivariate polynomials in species abundances. As mentioned in the previous section, to find the free-equilibrium points of system (4), we can set all time-derivatives to zero in order to obtain a system of multivariate polynomials in steady-state abundances 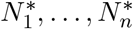 as follows

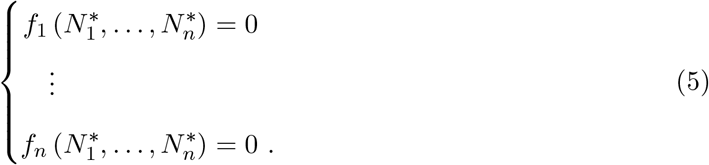

The number of free-equilibrium points of system (4) is given by the number of non-zero roots of system (5). While it is not a trivial problem, the number of non-zero roots can be calculated based on Bernshtein’s theorem (Bernshtein, 1975): Let us assume that the polynomial system has finitely many roots in (*C\**)^*n*^. Then the number of these roots is bounded from above by the mixed volume of its Newton polytopes *P*_*k*_, 1 ≤ *k* ≤ *n*. The upper bound of the number of non-zero roots is tight and achieved (exactly) for any generic choice of coefficients inside the polynomials *f*_1_, *f*_2_,…, *f*_*n*_ (note that in the LV model, when the vector of growth rates is not in the column space of the interaction matrix, there will be no solution and we can neglect such special cases). Therefore, Bernshtein’s theorem is the multivariate extension to the fact that a single variable polynomial of degree *n* will have *n*-complex roots for any generic coefficients.

To illustrate the quantification of non-zero roots of a polynomial system based on Bernshtein’s theorem, we show the details of how to compute Newton’s polytopes, the mixed volumes, and the evaluation of the number of complex roots using the following hypothetical system of equations (free-isoclines):

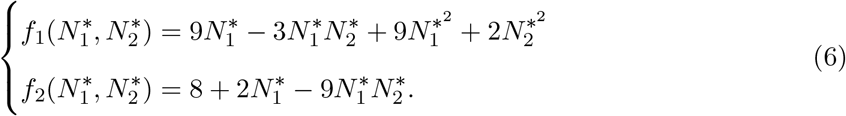

To compute the number of complex roots from Eqns. (6), we need to follow four basic concepts in algebraic geometry (Sommese & Wampler, 2005). For *i* = 1, 2:

1. We need to obtain the support *S*_*i*_ of *f*_*i*_, which is defined as the set of exponents of its monomials *eN*_*i*_’s. In system (6), the support *S*_1_ of *f*_1_ contains the points (*eN*_1_, *eN*_2_) in the set {(1,0),(1,1),(2,0),(0,2)}; while the support *S*_2_ of *f*_2_ contains the points (*eN*_1_, *eN*_2_) in the set {(0,0),(1,0),(1,1)}.
2. We need to obtain the Newton polytope *P*_*i*_ of *f*_*i*_, which is defined as the convex hull of the support *S*_*i*_. Fig. 1 (Panels A and B, respectively) shows the Newton polytopes *P*_1_ and *P*_2_ of system (6).
3. We need to perform the Minkowski sum *P*_*i*_ ⊕ *P*_*j*_ = {*p*_*i*_ + *p*_*j*_|*p*_*i*_ ∈ *P*_*i*_ and *p*_*j*_ ∈ *P*_*j*_} for *j* > *i*, which is defined as the convex hull of all possible summations of the supports *S*_*i*_ and *S*_*j*_. Fig. 1 (Panel C) shows *P*_1_ ⊕ *P*_2_ of system (6).
4. We need to obtain the mixed volume of the Newton polytopes *M* (*P*_1_, *P*_2_), which is defined as the difference in area between *P*_1_ ⊕ *P*_2_ and the sum of the areas of *P*_1_ and *P*_2_. The mixed volume corresponds to the exact number of roots that a multivariate polynomial system has. In system (6), the number of roots *M* (*P*_1_, *P*_2_) is given by

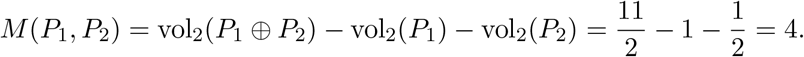

**Figure 1:**
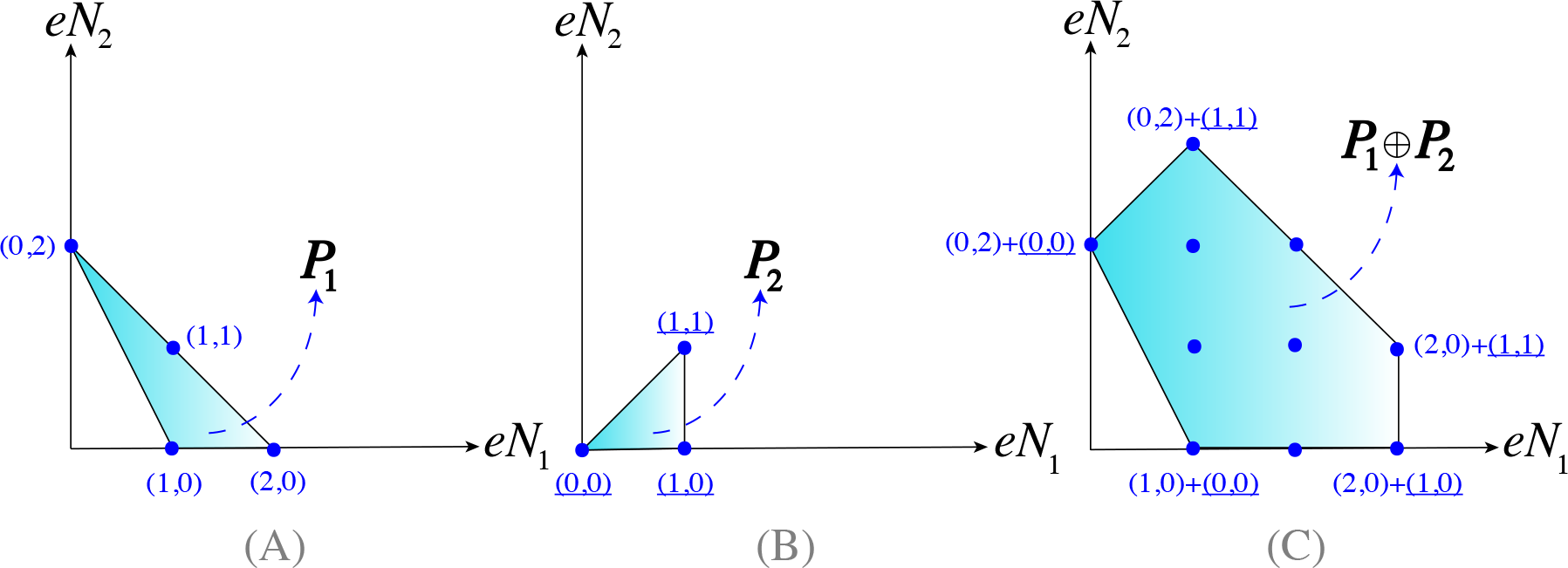
Illustration of the quantification of the number of complex roots of a polynomial system. For the hypothetical 2-species system defined by the set of Eqns. (6), the figure illustrates the construction of the mixed volume of Newton polytopes *P*_*i*_ of *f*_*i*_ for *i* = 1, 2. Panels **(A)** and **(B)** represent the Newton polytopes *P*_1_ and *P*_2_, respectively. Note that the coordinates (blue symbols) correspond to the different supports. Panel **(C)** represents the Minkowski sum of the first and second polytopes *P*_1_ ⊕ *P*_2_. Note that the mixed volume (number of complex roots) of this system is defined by *M* (*P*_1_, *P*_2_) = vol_2_(*P*_1_ ⊕ *P*_2_) − vol_2_(*P*_1_) − vol_2_(*P*_2_). The axes are the exponents of the supports’ monomials.

The calculation of the non-zero roots in the example above can be generalized to System (5) using the earlier steps and the formula of the mixed volume of *P*_1_,…, *P*_*n*_ (Sommese & Wampler, 2005) shown in Eqn. (7) below, which only requires computing the volumes of the Minkowski sums of all possible subsets of *P*_1_,…, *P*_*n*_ (see Gao et al. (2005); Lee & Li (2011) for methods and software packages to compute Eqn. (7) efficiently).

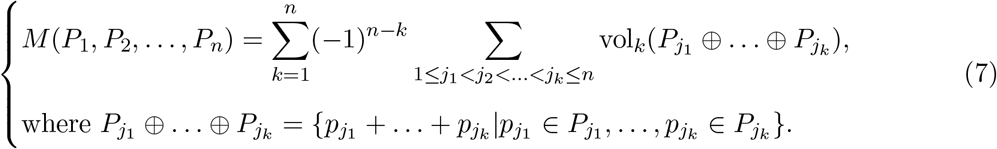

Note that if we remove the term 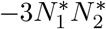 from 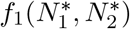 in Eqn. (6), then the number of non-zero roots of the system will not change. This is because the point (1, 1) in the support set *S*_1_ of *f*_1_ is not a corner point of the Newton polytope *P*_1_ (as it can be seen from Fig. 1A). Thus, removing that term (and parameter) will not affect the shape of *P*_1_. Importantly, this simple example illustrates that having more parameters in a multivariate polynomial system does not imply having more non-zero roots (i.e., more free-equilibrium points). Generally, all terms whose support coordinates are not corner points of the corresponding Newton polytope do not influence the number of non-zero roots in the multivariate polynomial system. This makes necessary to separate the problem of adding parameters to the problem of adding free-equilibrium points.

## Results

### Effects of HOIs on LV models

To investigate the difference in the number of free-equilibrium points between LV models with and without HOI terms, we followed our general methodology to calculate the number of non-zero roots in polynomial systems. In particular, we analytically computed the number of free-equilibrium points from the following 3 commonly used systems:

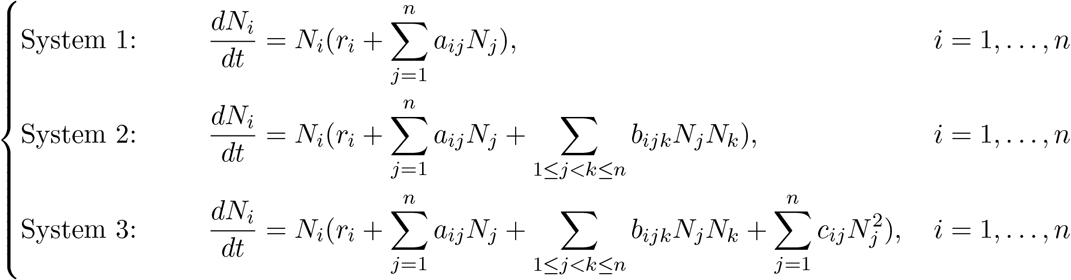

System 1 corresponds to the classic LV model without HOIs. While it is known that this system has only one free-equilibrium point (Takeuchi, 1996), using Bernshtein’s theorem (Bernshtein, 1975) we confirmed the existence of one single non-zero root. In turn, Systems 2 and 3 correspond to the simplest extensions of LV models with HOI terms. Note that System 3 has an additional higher-order self-regulation term. We found that Systems 2 and 3 have exactly 2^*n*^ − *n* and 2^*n*^ free-equilibrium points, respectively. Importantly, the increase in the number of parameters in a polynomial dynamical system does not imply an equal increase in the number of free-equilibrium points. For example, while System 3 has *n*^2^ terms more than System 2, it only has *n* free-equilibrium points more. Recall that only the corner terms to the corresponding Newton polytope determine the number of free-equilibrium points. In Systems 1 and 2, all the terms inside the brackets are corner terms. Similarly, in System 3, the *r′s* and the terms associated with the coefficients *c′s* are corner terms. However, the terms associated with the coefficients *a′s* and *b′s* are non-corner terms (see Supplementary Information for the mathematical derivations). This confirms that parameters and free-equilibrium points are two different descriptors of a dynamical model.

More generally, for the system

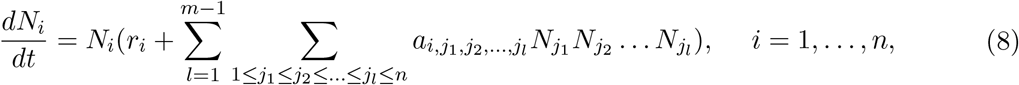

which represents *m*-order interactions in a LV model with HOI terms and *n* species, we found that the number of free-equilibrium points is given by (*m* − 1)^*n*^ (see Supplementary Information for the mathematical derivation). This result reveals that adding HOI terms to the LV model increases the number of free-equilibrium points exponentially with the dimension *n* of the system.

## Discussion

Recent work has shown that higher-order interactions can increase the stability (Grilli et al., 2017), promote the diversity (Bairey et al., 2016), and better explain the dynamics of ecological communities (Friedman et al., 2017; Mayfield & Stouffer, 2017). Yet, it has remained unclear whether the perceived explanatory power of adding higher-order terms into population dynamics models come from fundamental principles or a simple mathematical advantage given by the nature of multivariate polynomials. In fact, it has been perceived that mathematically there is nothing to prevent the inclusion of higher-order terms in ecological models (Letten & Stouffer, 2019).

Here, we have shown analytically that by adding HOI terms to ecological models, the number of free-equilibrium points increases exponentially with the dimension of the system. Recall that the classic LV model without HOI terms has a single free-equilibrium point, regardless of the number of parameters (Takeuchi, 1996). Importantly, we have shown that the more free-equilibrium points present in an ecological dynamical system, the more flexibility the system has to be engineered to fit experimental or observational data. This reveals that HOI terms cannot provide additional explanatory power of ecological dynamics if model parameters are not ecologically restricted.

The mathematical advantages coming from adding HOI terms into LV models can be easily seen in the mapping from the parameter space to the free-equilibrium point space of these systems (i.e., the steady-state species abundances). This mapping is one-to-one for the classic LV model, i.e., from ℝ^*θ*(*n*)^ (where *θ*(*n*) is the number of parameters in the model with *n* species) to ℝ^*n*^. However, when HOI terms are added to LV models, the mapping becomes from ℝ^*θ′*(*n*)^ to ℂ^*n*^, where *θ′*(*n*) *≥ θ*(*n*). Thus, the mapping becomes one to exponentially many when HOI terms are included. Also, the codomains for both mappings are different, it is ℝ^*n*^ for the classic LV model and ℂ^*n*^ for LV model with HOI terms. These explain why the number of both feasible and unfeasible solutions of a system increases when HOI terms are added. Note that even if studies penalize for the increase in the number of parameters *θ′*(*n*) − *θ*(*n*) between these models (e.g., using AIC, Akaike 1974), the mappings for HOI terms will continue to be from one to exponentially many, and the mathematical advantages will continue to be present. This reveals that models with and without HOI terms are fundamentally different and direct comparisons between them (e.g., dynamical properties or explanatory power) cannot be made.

To further illustrate the point above, one can focus on the problem of fitting data to ecological models. In this process, both parameters and initial conditions need to be tuned, introducing a bias. If a model has exponentially many free-equilibrium points, it is easier to engineer a feasible solution, especially without parameter restrictions. Thus fitting will be facilitated in models with higher-order terms (mainly due to the number of free-equilibrium points and not necessarily due to the number of parameters) and one may be tempted to conclude that feasibility is an ecological mechanism derived from higher-order interactions (while this is just a mathematical construct). Note that in this study, we have not started with a dynamical system and found how many of these free-equilibrium points are feasible (or stable). We have provided a general methodology to count the number of free-equilibrium points, which can be used as a measure of how easy it is to fit data to a dynamical model without parameter restrictions.

Finally, we would like to stress that if the aim of a study is to provide a comparison between models (e.g., proving a better fit or increasing conditions for stability and coexistence), then the two models should have at least a comparable number of free-equilibrium points under the same number of species. Otherwise, the mapping discussed earlier will be different for both models. Moreover, the parameter space for both models should be restricted to physical or ecological cases only (recall that by adding HOI terms it is possible to have imaginary equilibrium points but this is not possible in the classic LV model). Indeed, parameter constraints can help us to quantify more fairly the parameter space compatible with critical ecological behavior between models (Saavedra et al., 2017; Song et al., 2018). Because explaining and predicting are two different tasks (Shmueli, 2010), here we advise that ecological models with higher-order terms may be good for prediction purposes, but their explanatory power should be taken with caution.

## Acknowledgments

MA would like to thank Kuwait Foundation for the Advancement of Sciences (KFAS) and Kuwait Chamber of Commerce and Industry (KCCI) for supporting and funding his PhD studies. Funding was provided by the Mitsui Chair to SS. We would also like to thank Simone Cenci, Lucas Medeiros, Rudolf P. Rohr, and Chuliang Song for fruitful discussions about this manuscript.

## Author contributions

SS and MA designed the study, MA performed the study, SS supervised the study, SS and MA analyzed results and wrote the manuscript.

## Supplementary Information

### Derivation of the number of free- and total-equilibrium points

In the Results section of the main text, we presented the findings of the number of free-equilibrium points in three different LV models (one with and two without HOI terms). Here, we present the formal proof of their derivation. The proof is divided into two parts. Free-equilibrium points are considered in Part 1 while the total number of points are considered in Part 2 (free + rigid) for the following studied systems

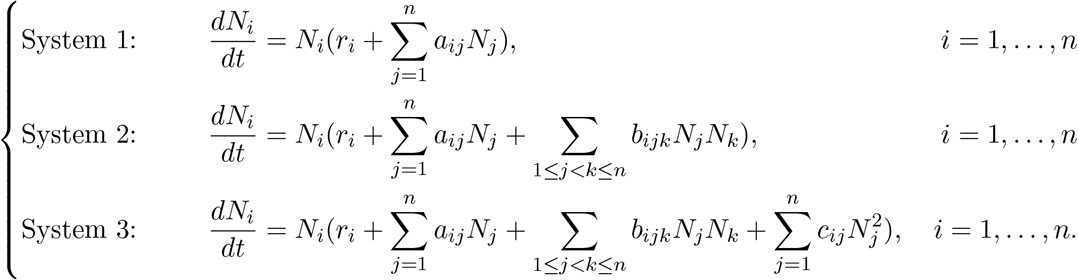

#### Part 1

To find the free-equilibrium points, we set all time-derivatives to zero in the systems above. We ignore the *N*_*i*_ terms outside the brackets because they produce solutions which have at least one zero component (rigid-equilibrium points) which will be considered in Part 2 of this section. Therefore, the equations we need to study are

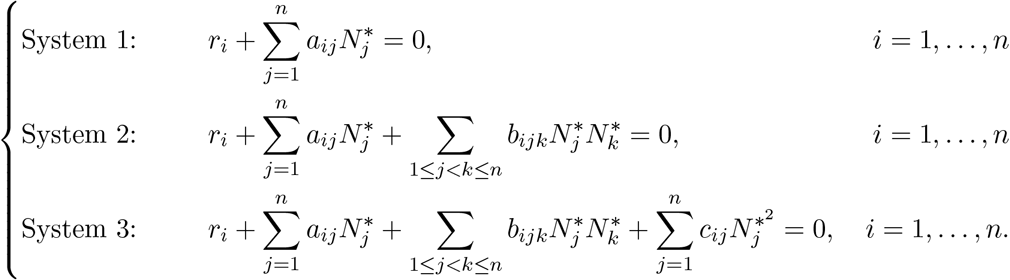

Note that all the equations for each of the 3 systems are functions of the species abundances and contain the exact same terms. Hence, the support of each equation, and thus the Newton polytopes, are identical and we will denote them by *S*_(1)_ and *P*(*S*_(1)_), respectively. Let *S*_(*k*)_ be defined as 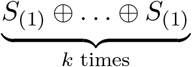. Furthermore, let us define *e*_*i*_ to be the point in space with its *i*^th^ component to be 1 and the rest are all zeros. It is also important to note that the operations of the Minkowski summation and those of forming convex hulls are commuting (Abardia-Evquoz et al., 2018), that is

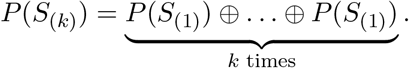

#### System 1

Focusing on System 1, it is easy to see that the vertices of *P*(*S*_(1)_) are the origin and *e*_*i*_ for *i* = 1,…, *n*. Note that all the terms in System 1 are corner points. To perform the induction step, let us assume that the vertices of *P*(*S*_(*k*)_) are the origin and *ke*_*i*_ for *i* = 1,…, *n*. *P*(*S*_(*k*+1)_) = *P*(*S*_(*k*))_)⊕*P*(*S*_(1)_) imply that the vertices of *P*(*S*_(*k*+1)_) are contained in the Minkowski sum of the verticies of *P*(*S*_(*k*))_) and *P*(*S*_(1))_), which are the origin, *e*_*i*_, *ke*_*i*_ and *ke*_*i*_ + *e*_*j*_ for *i*, *j* = 1,…, *n*. It is useful to isolate the case *i* = *j* from the term *ke*_*i*_ + *e*_*j*_ to the standalone term (*k* + 1)*e*_*i*_. Hence, the Minkowski sum of the verticies of *P*(*S*_(*k*))_) and *P*(*S*_(1))_) are the origin, *e*_*i*_, *ke*_*i*_, *ke*_*i*_ + *e*_*j*_ and (*k* + 1)*e*_*i*_ for *i*, *j* = 1,…, *n* and *i* ≠ *j*. Note that both *e*_*i*_ and *ke*_*i*_ lie in the line connecting the origin and (*k* + 1)*e*_*i*_ for *i* = 1,…, *n*, hence, they cannot be vertices of *P*(*S*_(*k*+1)_). Moreover, *ke*_*i*_ + *e*_*j*_ lie in the line connecting (*k* + 1)*e*_*i*_ and (*k* + 1)*e*_*j*_ for *i*, *j* = 1,…, *n* and *i* ≠ *j*. Hence, the vertices of *P*(*S*_(*k*+1)_) are the origin and (*k* + 1)*e*_*i*_ for *i* = 1,…, *n*. Thus, induction is complete. Therefore for all, positive integers *k*, the vertices of *P*(*S*_(*k*)_) are the origin and *ke*_*i*_ for *i* = 1,…, *n*.

The computation of the volume of *P*(*S*_(*k*)_) is simply equivalent to finding the volume of the generalized-tetrahedron that is bounded by the coordinate hyperplanes and the hyperplane *eN*_1_ + ⋯ + *eN*_*n*_ = *k*, which is

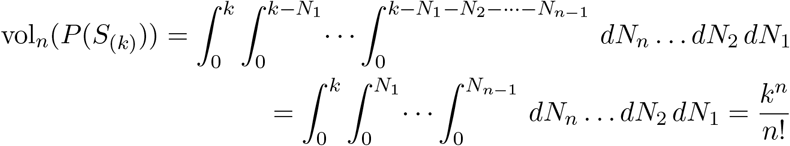

under a change of variables to the upper limits of the integrands. To find the number of non-zero roots, we just need to compute the mixed volume, which is

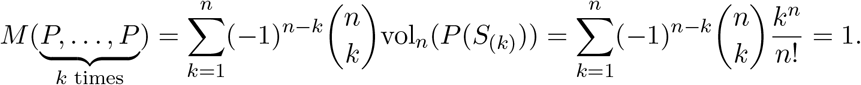

Thus, we have provided an alternative proof to confirm that the classic LV model has only 1 free-equilibrium point.

**Table S1:** The table shows the Minkowski sum between the vertices of *P*(*S*_(*k*)_) (which are the origin, *ke*_*i*_, and *ke*_*i*_ + *ke*_*j*_) and the vertices of *P*(*S*_(1)_) (which are the origin, *e*_*i*_, and *e*_*i*_ + *e*_*j*_ for *i*, *j*, *l*, *m* = 1,…, *n* where *i* ≠ *j* ≠ *l* ≠ *m*). All single bracketed terms are the result of the Minkowski sum of these terms with the origin. Double bracketed terms are the Minkowski sum of the expression in the first bracket with that in the second one. All coordinates which contain (*k* + 1) are special cases of the latter double bracketed expression in the table when a term in the second bracket combines with one in the first bracket. Note that in the 8^th^ line (*k* + 1)*e*_*i*_ + *e*_*j*_ is mentioned without any mentioning to *e*_*i*_ + (*k* + 1)*e*_*j*_ as the entire set of coordinates generated by both expressions are identical for *i*, *j* = 1,…, *n* and *i* ≠ *j*.

| Coordinate | Start point | End point |
| --- | --- | --- |
| [0] | - | - |
| $[e_i]$ | 0 | $(k+1)e_i$ |
| $[e_i + e_j]$ | 0 | $(k+1)e_i + (k+1)e_j$ |
| $[ke_i]$ | 0 | $(k+1)e_i$ |
| $[ke_i] + [e_j]$ | $(k+1)e_i$ | $(k+1)e_j$ |
| $(k+1)e_i$ | - | - |
| $[e_i + e_j] + [ke_l]$ | $(k+1)e_i + (k+1)e_j$ | $(k+1)e_l$ |
| $(k+1)e_i + e_j$ | $(k+1)e_i$ | $(k+1)e_i + (k+1)e_j$ |
| $[ke_i + ke_j]$ | 0 | $(k+1)e_i + (k+1)e_j$ |
| $[ke_i + ke_j] + [e_l]$ | $(k+1)e_i + (k+1)e_j$ | $(k+1)e_l$ |
| $(k+1)e_i + ke_j$ | $(k+1)e_i$ | $(k+1)e_i + (k+1)e_j$ |
| $[ke_i + ke_j] + [e_l + e_m]$ | $(k+1)e_i + (k+1)e_j$ | $(k+1)e_l + (k+1)e_m$ |
| $(k+1)e_i + ke_j + e_l$ | $(k+1)e_i + (k+1)e_j$ | $(k+1)e_i + (k+1)e_l$ |
| $(k+1)e_i + (k+1)e_j$ | - | - |

##### System 2

Focusing on System 2, the vertices of *P*(*S*_(1)_) are the origin, *e*_*i*_, and *e*_*i*_ + *e*_*j*_ for *i*, *j* = 1,…, *n* and *i* < *j*. Note that all the terms in System 2 are corner points. To perform an induction step, let us assume that the vertices of *P*(*S*_(*k*)_) are the origin, *ke*_*i*_, and *ke*_*i*_ + *ke*_*j*_ for *i*, *j* = 1,…, *n* and *i* < *j*. Again, since *P*(*S*_(*k*+1)_) = *P*(*S*_(*k*))_) ⊕ *P*(*S*_(1)_), then the vertices of *P*(*S*_(*k*+1)_) are contained in the Minkowski sum of the verticies of *P*(*S*_(*k*))_) and *P*(*S*_(1))_), which are shown in Table S1. Note that for the points that are not corner points, their starting and ending points are also included in the table. From Table S1, it is easy to see that the vertices of *P*(*S*_(*k*+1)_) are the origin, (*k* + 1)*e*_*i*_ and (*k* + 1)*e*_*i*_ + (*k* + 1)*e*_*j*_ for *i*, *j* = 1,…, *n* and *i* < *j*. Thus, induction is complete. Therefore, for all positive integers *k*, the vertices of *P*(*S*_(*k*)_) are the origin, *ke*_*i*_, and *ke*_*i*_ + *ke*_*j*_ for *i*, *j* = 1,…, *n* and *i* < *j*.

To compute the mixed volume (hence the number of non-zero roots), let us define 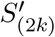 as the union of the verticies of *P*(*S*_(*k*)_) as well as 2*ke*_*i*_ for *i* = 1,…, *n*. Note that the vertices of 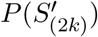 are simply the origin and 2*ke*_*i*_ for *i* = 1,…, *n*. This is true given that *ke*_*i*_ lies in the interior of the line connecting the origin and 2*ke*_*i*_, whereas *ke*_*i*_ + *ke*_*j*_ lies in the line connecting 2*ke*_*i*_ and 2*ke*_*j*_ for *i*, *j* = 1,…, *n* and *i* < *j*. Also note that *P*(*S*_(*k*)_) is contained in 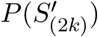 and the difference in their volumes is the sum of the individual volumes of the generalized-tetrahedron_*i*_ for *i* = 1,…, *n*—whose vertices are *ke*_*i*_, *ke*_*i*_ + *ke*_*j*_, and 2*ke*_*i*_ (we just exclude the origin) for *j* = 1,…, *n* and *j* ≠ *i* (see Figure S1). These volumes are all identical and there are *n* of them. To help to visualize this, we can shift each of these coordinates of the generalized-tetrahedron_*i*_ by *ke*_*i*_ to get the shifted structure, whose vertices are the origin and *ke*_*i*_ for *i* = 1,…, *n*—which is exactly 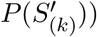. Thus, the volume of *P*(*S*_(*k*)_) is

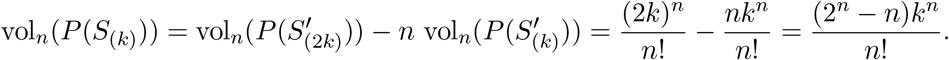

To find the number of non-zero roots, we just need to compute the mixed volume which is

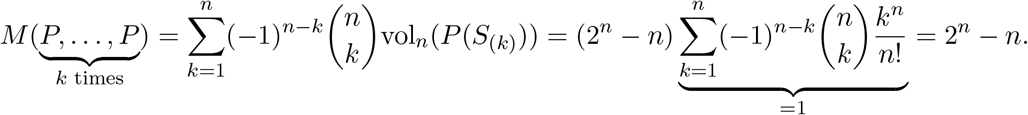

**Figure S1:**
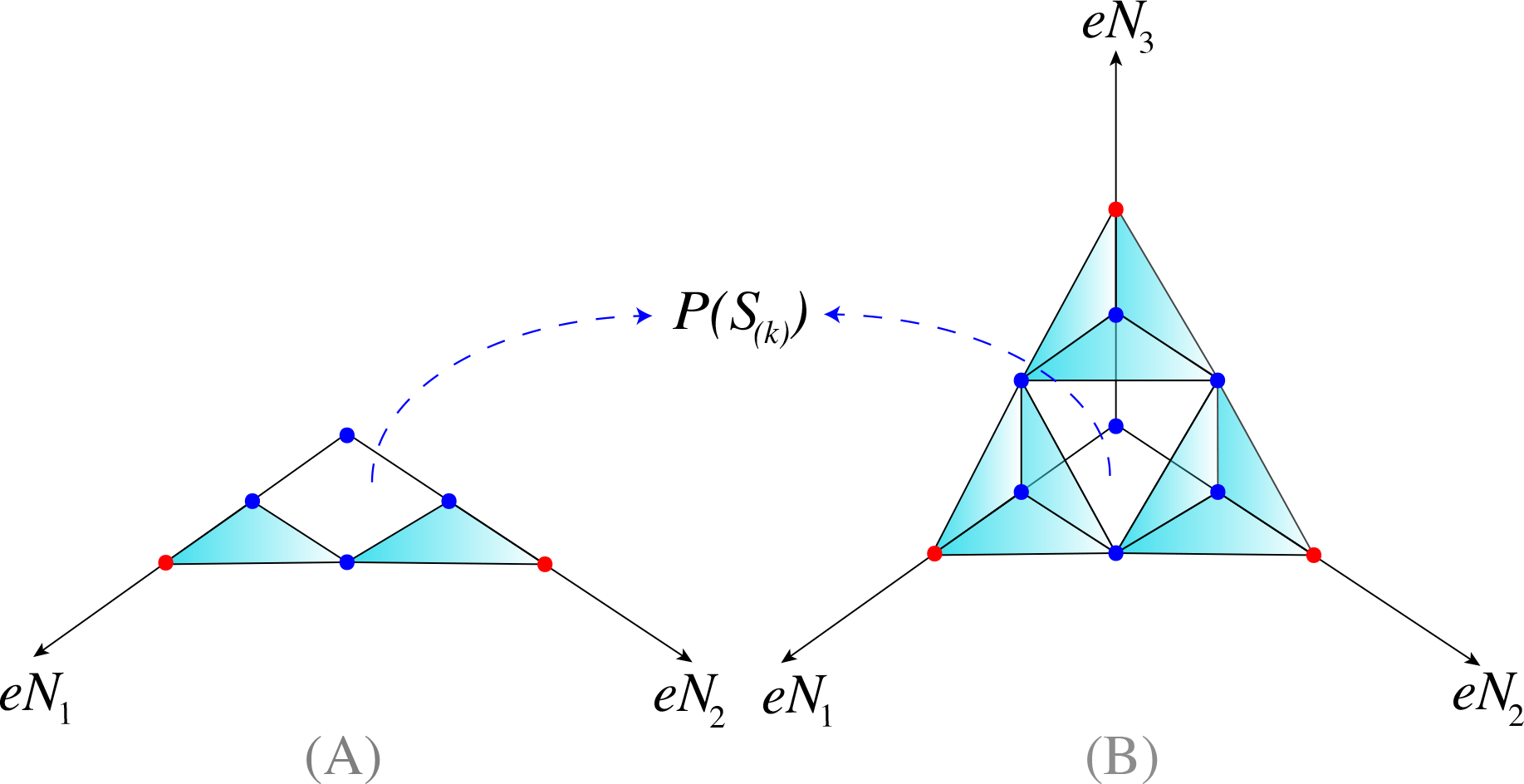
Illustration of the difference in volumes between 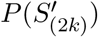 (which are the cyan plus white regions) and *P*(*S*_(*k*)_) (which are the white regions) for two and three species equations is shown in panels **(A)** and **(B)** respectively. The blue dots are vertices of *P*(*S*_(*k*)_), which are spaced uniformly by *k* units along the axes, while the red and the origin are vertices of 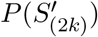. The axes are the exponents of the supports’ monomials.

##### System 3

Focusing on System 3, the support *S*_(1)_ contains the origin, *e*_*i*_, *e*_*i*_ + *e*_*j*_, and 2*e*_*i*_ for *i*, *j* = 1,…, *n* and *i* < *j*. Therefore, the vertices of *P*(*S*_(1)_) are the origin and 2*e*_*i*_ for *i* = 1,…, *n*. This is true given that *e*_*i*_ and *e*_*i*_ + *e*_*j*_ are not corner points of *P*(*S*_(1)_) for *i*, *j* = 1,…, *n* and *i* < *j*. That is in System 3, the *r′s* and the terms associated with the coefficients *c′s* are corner terms. However, the terms associated with the coefficients *a′s* and *b′s* are non-corner terms. To perform an induction step, let us assume that the vertices of *P*(*S*_(*k*)_) are the origin and 2*ke*_*i*_ for *i* = 1,…, *n*. *P*(*S*_(*k*+1)_) = *P*(*S*_(*k*))_) ⊕ *P*(*S*_(1)_) imply that the vertices of *P*(*S*_(*k*+1)_) are contained in the Minkowski sum of the verticies of *P*(*S*_(*k*))_) and *P*(*S*_(1))_)—which are the origin, 2*e*_*i*_, 2*ke*_*i*_ and 2*ke*_*i*_ + 2*e*_*j*_ for *i*, *j* = 1,…, *n*. It is useful to isolate the case *i* = *j* from the term 2*ke*_*i*_ + 2*e*_*j*_ to the standalone term 2(*k* + 1)*e*_*i*_. Hence, the Minkowski sum of the verticies of *P*(*S*_(*k*))_) and *P*(*S*_(1))_) are the origin, 2*e*_*i*_, 2*ke*_*i*_, 2*ke*_*i*_ + 2*e*_*j*_, and 2(*k* + 1)*e*_*i*_ for *i*, *j* = 1,…, *n* and *i* ≠ *j*. Note that both 2*e*_*i*_ and 2*ke*_*i*_ lie in the line connecting the origin and 2(*k* + 1)*e*_*i*_ for *i* = 1,…, *n*, hence, they cannot be vertices of *P*(*S*_(*k*+1)_). Moreover, 2*ke*_*i*_ + 2*e*_*j*_ lies in the line connecting 2(*k* + 1)*e*_*i*_ and 2(*k* + 1)*e*_*j*_ for *i*, *j* = 1,…, *n* and *i* ≠ *j*. Hence, the vertices of *P*(*S*_(*k*+1)_) are the origin and 2(*k* + 1)*e*_*i*_ for *i* = 1,…, *n*. Thus, induction is complete. Therefore, for all positive integers *k*, the vertices of *P*(*S*_(*k*)_) are the origin and 2*ke*_*i*_ for *i* = 1,…, *n*. From the derivation in System 1, we already know that the volume of *P*(*S*_(*k*)_) is (2*k*)^*n*^/*n*! (it is the volume of the generalized-tetrahedron that is bounded by the coordinate hyperplanes and the hyperplane *eN*_1_ + ⋯ + *eN*_*n*_ = 2*k*). Then, to find the number of non-zero roots, we just need to compute the mixed volume which is

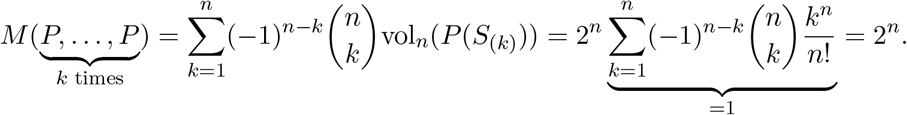

#### Part 2

Here, we calculate *M*_*T*_—the total number of equilibrium-points for any of the 3 systems under investigation, including their rigid-equilibrium points (the ones which have one or more zero-components). Note that rigid-equilibrium points of LV model with or without HOI terms are the free-equilibrium points of the same model after substituting the corresponding zero abundances in them and deleting the lines which involve their time derivative as they will be zero as well. To see this, we can always start with an LV model with or without higher-order interaction terms and focus on the rigid-equilibrium point *O*_*m*_ in which species *m* dies out. That is, *N*_*m*_ = 0 in *O_m_*. Note that these models do not allow revival of species. Therefore, when a species dies out, the value *N*_*m*_ = 0 will stay the same forever (i.e., *dN*_*m*_/*dt* = 0). Upon substituting *N*_*m*_ = 0 into the original dynamical system and deleting the line *dN*_*m*_/*dt* = 0 from it, we get a model which is one species less for which *O*_*m*_ but with the point/coordinate *N*_*m*_ = 0 is deleted from it to be its free-equilibrium point. The remaining *n* − 1 equations will have identical polytopes but with all the terms involving *N*_*m*_ being removed. Therefore, the new systems will have exactly *M*_*n*−1_ free-equilibrium points, providing *n* ways to eliminate a single species. Under the same logic, by letting *k* species go extinct, the new systems will have *M*_*n*−*k*_ free-equilibrium points with *nC*_*k*_ ways to do it. Hence,

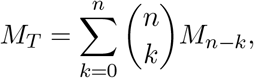

Recall that from Part 1 we already know that for *k* species, the number of free-equilibrium points for systems 1, 2, and 3 are 1, 2^*k*^ − *k*, and 2^*k*^, respectively. Therefore,

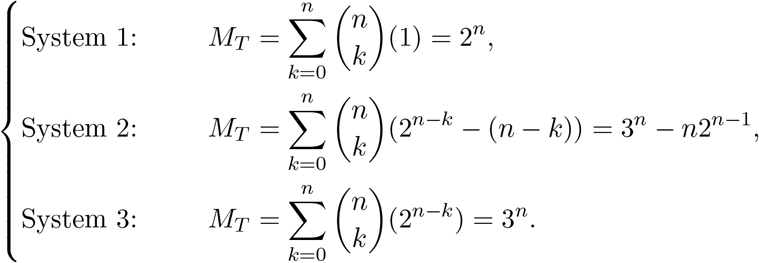

The results above confirm that the LV model has a total of 2^*n*^ equilibrium points (i.e., free-equilibrium points + rigid-equilibrium points) (Takeuchi, 1996). Importantly, we can also clearly see that adding higher-order terms makes the total number of equilibrium points jump from 2^*n*^ to 3^*n*^ − *n*2^*n*−1^ when we add non-additive pairwise interactions terms per capita (System 2), furthermore, these number jumps to 3^*n*^ when we include the non-additive quadratic terms per capita (System 3).

### Derivation of the number of free- and total-equilibrium points for a general LV system with HOIs

For the general system shown below:

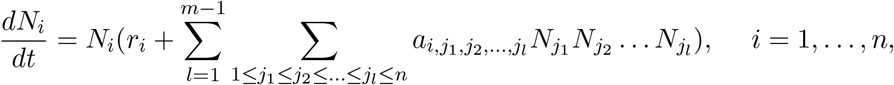

The support *S*_(1)_ contains the origin, 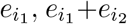,… and 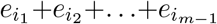 for *i*_1_*, i*_2_,…, *i*_*m*−1_ = 1, 2,…, *n*. All these coordinates are bounded by the coordinate hyperplanes and the hyperplane *eN*_1_ +… + *eN*_*n*_ = (*m* − 1). Hence, the origin and (*m* − 1)*e*_*i*_ for *i* = 1,…, *n* are the vertices of *P*(*S*_(1)_) where the term (*m* − 1)*e*_*i*_ is obtained by setting *i*_1_ = *i*_2_ =… = *i*_*m*−1_ ≡ *i* in 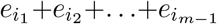. To perform an induction step, let us assume that the vertices of *P*(*S*_(*k*)_) are the origin and 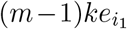 for *i* = 1,…,*n*. Since *P*(*S*_(*k*+1)_) = *P*(*S*_(*k*))_)⊕*P*(*S*_(1)_), then the vertices of *P*(*S*_(*k*+1)_) are contained in the Minkowski sum of the verticies of *P*(*S*_(*k*))_) and *P*(*S*_(1))_)—which are the origin, (*m* − 1)*e*_*i*_, (*m* − 1)*ke*_*i*_ and (*m* − 1)*ke*_*i*_ + (*m* − 1)*e*_*j*_ for *i*, *j* = 1,…, *n*. It is useful to isolate the case *i* = *j* from the term (*m*− 1)*ke*_*i*_ +(*m*− 1)*e*_*j*_ to the standalone term (*m*− 1)(*k* +1)*e*_*i*_. Therefore, the vertices of *P*(*S*_(*k*+1)_) are the origin, (*m* − 1)*e*_*i*_, (*m* − 1)*ke*_*i*_, (*m* − 1)*ke*_*i*_ + (*m* − 1)*e*_*j*_, and (*m* − 1)(*k* + 1)*e*_*i*_ for *i*, *j* = 1,…, *n* and *i* ≠ *j*. Note that both (*m* − 1)*e*_*i*_ and (*m* − 1)*ke*_*i*_ lie in the line connecting the origin and (*m*− 1)(*k* + 1)*e*_*i*_ for *i* = 1,…, *n*, hence, they cannot be vertices of *P*(*S*_(*k*+1)_). Moreover, (*m* − 1)*ke*_*i*_ + (*m* − 1)*e*_*j*_ lies in the line connecting (*m* − 1)(*k* + 1)*e*_*i*_ and (*m* − 1)(*k* + 1)*e*_*j*_ for *i*, *j* = 1,…, *n* and *i* ≠ *j*. Hence, the vertices of *P*(*S*_(*k*+1)_) are the origin and (*m* − 1)(*k* + 1)*e*_*i*_ for *i* = 1,…, *n*. Thus, induction is complete. Therefore, for all positive integers *k*, the vertices of *P*(*S*_(*k*)_) are the origin and (*m* − 1)*ke*_*i*_ for *i* = 1,…, *n*. From the derivation in System 1, we already know that the volume of *P*(*S*_(*k*)_) is ((*m* − 1)*k*)^*n*^/*n*! (it is the volume of the generalized-tetrahedron that is bounded by the coordinate hyperplanes and the hyperplane *eN*_1_ +…+*eN*_*n*_ = (*m*−1)*k*). Then, to find the number of non-zero roots, we just need to compute the mixed volume which is

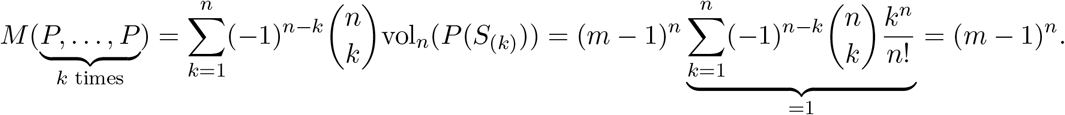

Unlike corner terms, if non-corner terms are removed from the generalized model (non-corner points of the generalized-tetrahedron that is bounded by the coordinate hyperplanes and the hyperplane *eN*_1_ + ⋯ + *eN*_*n*_ = *m* − 1), the number of free-equilibrium points will not be affected. However, removing corner points will reduce that number as we have seen in System 2 which is essentially System 3 but with some of its corner points being removed. Regards to the number of total (rigid+free) equilibrium points for this general system, it is given by

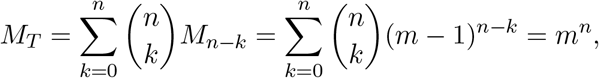

